# Detection of tomato brown rugose fruit virus in environmental residues: the importance of contextualizing test results

**DOI:** 10.1101/2024.04.25.591117

**Authors:** Anne K.J. Giesbers, Elise Vogel, Anna Skelton, Zafeiro Zisi, Mandy Wildhagen, Yue L. Loh, Lucas Ghijselings, Johanne Groothuismink, Marcel Westenberg, Jelle Matthijnssens, Annelien Roenhorst, Christine Vos, Adrian Fox, Marleen Botermans

## Abstract

Tomato brown rugose fruit virus (ToBRFV) is regulated as a quarantine pest in many countries worldwide. To assess whether ToBRFV is present in cultivations, plants or seed lots, testing is required. The interpretation of test results, however, can be challenging. Real-time RT-PCR results, even though considered “positive”, may not always signify plant infection or indicate the presence of infectious virus, but could be due to the presence of viral residues in the environment. Here, case studies from the Netherlands, Belgium, and the United Kingdom address questions regarding the detection of ToBRFV in various settings, and the infectiousness of ToBRFV positive samples. These exploratory analyses demonstrate widespread detection of ToBRFV in diverse samples and environments. ToBRFV was detected inside and around greenhouses with no prior history of ToBRFV infection, on different materials and surfaces including those that were untouched by individuals, plants, or objects. This suggested the dispersal of viral residues through aerosols. ToBRFV or its residues were more often detected in areas with nearby tomato production yet were also found in a wider environment extending beyond infected crops. Given that ToBRFV originating from environmental contamination may or may not be infectious, adds complexity to decision-making in response to positive test results. Contextual information, such as the origin of the sample and the likelihood of residues from prior cultivations and/or the broader environment, is important for interpreting test results. A nuanced approach is crucial to correctly interpret ToBRFV test results, necessitating further research to support risk assessment.

## Introduction

Tomato brown rugose fruit virus (ToBRFV) is a tobamovirus that may cause severe disease in tomato and pepper crops. Therefore, it is regulated as quarantine pest in numerous countries, necessitating testing to establish its presence or absence (EPPO, 2024). ToBRFV was initially identified in 2015 in Jordan (Salem *et al*., 2016), and has currently been reported from Europe, Africa, North America, South America and Asia (EPPO, 2024). Natural hosts of ToBRFV include tomato (*Solanum lycopersicum*) and pepper (*Capsicum* spp.) (Luria *et al*., 2017, Panno *et al*., 2020, Salem *et al*., 2020). Infected tomato plants may show mosaic patterns on leaves, narrowing of leaves, and the appearance of yellow spots or brown wrinkled (rugose) symptoms on fruits (Luria et al., 2017, Salem et al., 2016). ToBRFV poses a significant threat to tomato production, given that the established tobamovirus resistance genes in tomatoes, namely *Tm-1*, *Tm-2* and *Tm-2^2^*, are ineffective against this virus (Luria et al., 2017). For pepper, reported symptoms encompass yellowing, necrotic lesions or spots, and discoloration of fruits. The resistance provided by the *L^3^*and *L^4^* genes or alleles may break down at temperatures exceeding 30°C (Eldan *et al*., 2022, Fidan *et al*., 2021).

Like other tobamoviruses, ToBRFV is highly stable and easily transmitted through contact and propagation materials. Furthermore, ToBRFV is seedborne and seed-transmitted, as it can be located externally on the seed coat (Davino *et al*., 2020, Salem *et al*., 2022b). Several territories have listed ToBRFV as a quarantine pest and the European Union (EU) has implemented emergency measures since 2019 for tomato and pepper (EU, 2019), to prevent the entry and spread of the virus. Phytosanitary controls within these emergency measures include statutory surveillance through testing by (real-time) RT-PCR. In addition, appropriate measures to eradicate ToBRFV from infected production sites are required by the end of the cropping cycle.

However, it has been hypothesized that positive test results do not always indicate plant infection but could be attributed to the presence of residual virus in the environment, for instance originating from highly infected plants in close proximity (Skelton *et al*., 2023b). This raises questions regarding how to correctly interpret positive test results and the potential risks associated with a broader presence of the virus in the environment. The virus’s persistent nature, the ease of transmission through contact and presumed environmental spread coupled with highly sensitive diagnostic tests, make interpreting positive test results complex. It is particularly challenging to discern whether plants are genuinely infected or if the detected presence of ToBRFV is due to environmental contamination.

Assessment of the phytosanitary risk posed by environmental residues adds another layer of complexity. There has been limited work related to the phytosanitary relevance and transmission risk of environmental ToBRFV. One study indicated that irrigation with ToBRFV-contaminated water can lead to plant infection (Mehle *et al*., 2023). Another study suggested that the presence of ToBRFV residues in a greenhouse could lead to crop infection. The latter hypothesis was supported by the identification of almost identical ToBRFV sequences in a tomato crop during a ToBRFV outbreak and in the subsequent crop in the same greenhouse several months after disinfection and crop rotation (van de Vossenberg *et al*., 2021). Apart from the transmission risk, ToBRFV in environmental residues also poses a ‘diagnostic risk’ of erroneously reporting crop infection in the absence of truly infected plants (Skelton et al., 2023b).

To gain more insight into the detection of ToBRFV in various environmental settings and its potential phytosanitary implications, case studies were conducted in the Netherlands, Belgium, and the United Kingdom (UK). Collectively, these studies focused on the following questions:

1. Can ToBRFV be detected on presumed nonhost plants and inanimate surfaces, not in direct contact with infected plants, individuals, or objects, suggesting environmental contamination (ToBRFV residues)? (case study 1, 2, 3, 5)
2. Can ToBRFV residue be infectious? (case study 1, 3, 4, 6)
3. Is there a distinction between locations with and without prior ToBRFV outbreaks? (case study 1)
4. Are there differences between regions with high and low concentrations of (infected) tomato production? (case study 1)

These exploratory analyses demonstrate the presence of ToBRFV residues in diverse settings, even in the absence of direct contact with infection sources, thereby offering insights into the potential implications caused by the presence of environmental residues.

## Materials and Methods

### Case study 1 (Netherlands)

Various samples, including leaves, swabs from a range of surfaces, water and soil or substrate, were collected in and around greenhouses in the Netherlands producing either cucumber (*Cucumis sativus*) or green bean (*Phaseolus vulgaris*), which are nonhosts of ToBRFV. Some of the greenhouses had grown tomato and experienced a ToBRFV outbreak in the past half year before sampling, but after crop cleanout and disinfection they had switched to the production of a nonhost crop in all or part of their compartments. The geographic regions where sampling was carried out either had a high or a low concentration of tomato production in the West of the Netherlands (sampled in June-July 2023) and the East of the Netherlands (sampled in October 2023) respectively (Fig 1a). The choice of samples per location differed slightly (Table 1), due to the exploratory nature of this study and the unique characteristics of each location. The primary sample types consisted of swabs and leaves. Swabs were collected from surfaces that were either typically untouched (e.g. ceiling structures) or had been exposed to contact with individuals, plants or objects. For both cucumber and green bean, lower leaves were sampled, along with dead leaves from the floor. For tomato, young leaves from the top of the plants were bulk sampled. Various additional samples included soil from beneath the plastic covers, rockwool, paperwork, volunteer tomato plants (company 6 only), wastewater, rainwater, and disinfected water. Outside greenhouses, leaf and swab samples were collected from various plants and different surfaces.

**Figure 1.**
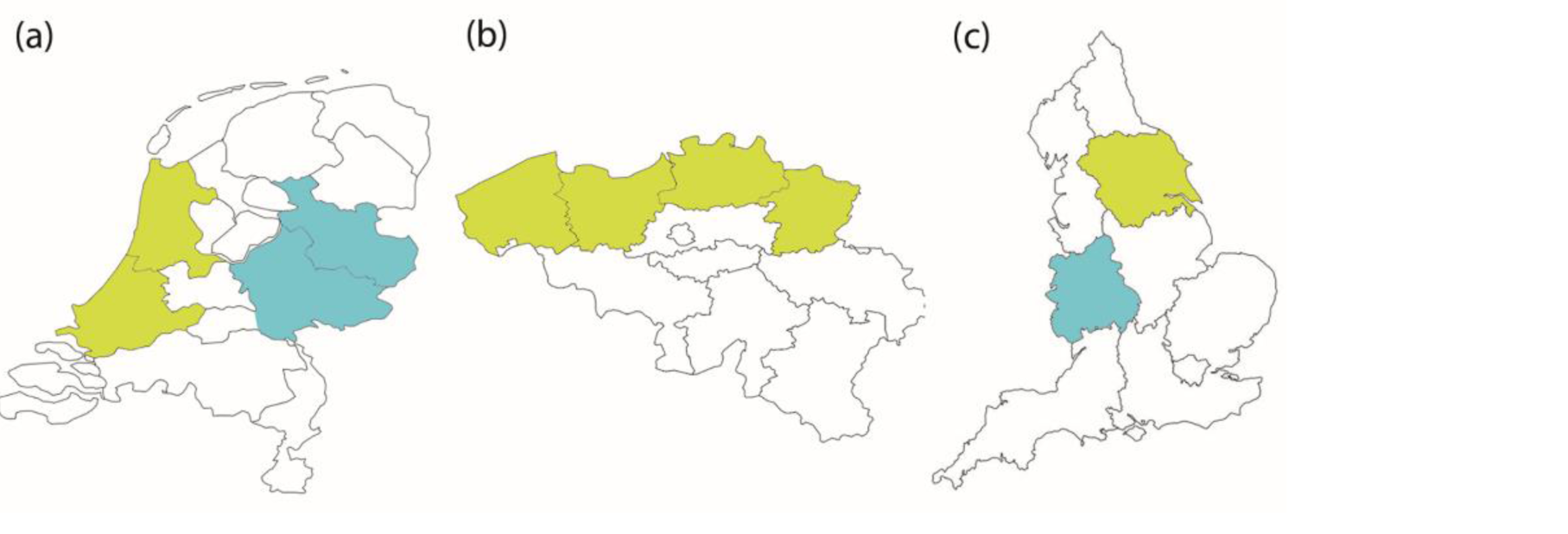
Regions of sampling a) West of the Netherlands (yellow; case study 1 and 2), East of the Netherlands (blue; case study 1) b) North of Belgium (yellow; case study 3 and 4) c) West Midlands, UK (blue; case study 5), North-East of England, UK (yellow; case study 6). Maps are not drawn to scale. Companies/locations were unique for each case study, exact geographic locations were omitted for confidentiality reasons.

**Table 1.**
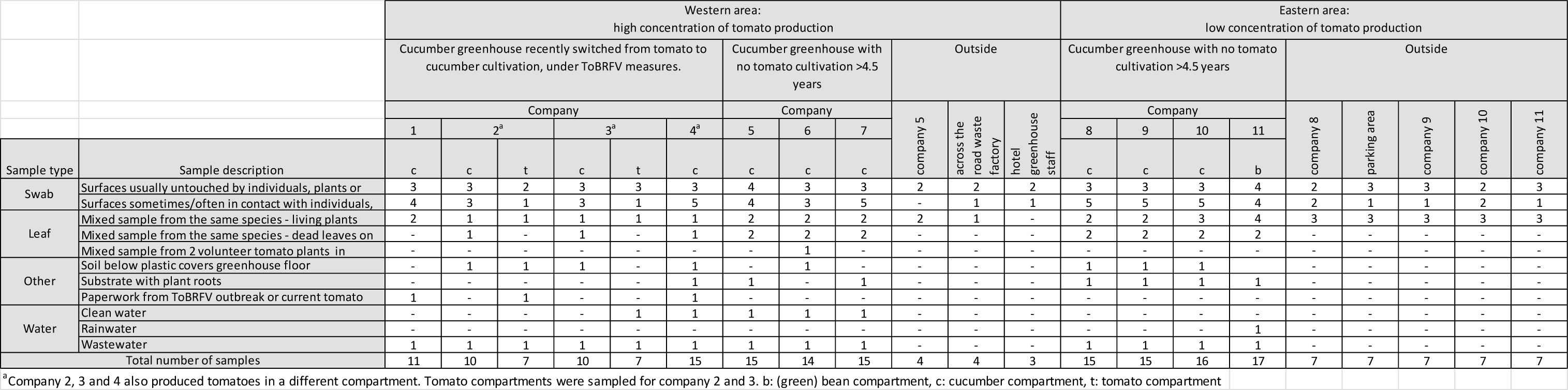
Number and types of samples per location and area.

### Case study 2 (Netherlands)

Leaf or sepal samples of tomato (*S. lycopersicum*), drain water and swab samples from different surfaces with varying levels of contact with individuals, plants, or objects, were collected at three adjoining tomato production greenhouses belonging to the same company in the West of the Netherlands (Fig 1a). To our knowledge, none of the sampled locations had been previously infected with ToBRFV. Samples were collected in the summer of 2021.

### Case study 3 (Belgium)

Weed samples were collected within 3 m outside of a tomato production greenhouse situated in an area with a high concentration of tomato production in the North of Belgium in the summer of 2023 (Fig 1b). This company likely experienced a ToBRFV outbreak at the time of sampling, as the National Plant Protection Organization had been notified of an infection earlier in the season. The following weed species, common to Western Europe, were sampled randomly: *Erigeron canadensis, Galinsoga quadriradiata, Solanum nigrum,* and *Sonchus oleraceus*.

### Case study 4 (Belgium)

High throughput genome sequencing (HTS) analysis was performed on two tomato leaf samples collected in 2020-2021 from two adjoining tomato production greenhouses of the same company situated in an area with a high concentration of tomato production in the North of Belgium (Fig 1b). The crops at both locations became infected with ToBRFV shortly after each other.

### Case study 5 (United Kingdom)

In February 2022, swab samples were collected from different surfaces in a tomato production facility consisting of two conjoined greenhouses, in the West Midlands, UK (Fig 1c) with an ongoing ToBRFV outbreak, first detected in March 2020. Many of the sampled surfaces had not been in direct contact with individuals, plants, or objects, e.g., the greenhouse roof structure (inside), tops of roof lights, and inside fans. Other surfaces included bee boxes, the underside of the gutters, a Tyvek suit, and the grower’s phone and gloves. The Tyvek suit was sampled immediately after it was removed by the person taking the swab samples. The sampled area specifically targeted the section where the person had leaned against a retaining bar on a scissor lift, for support, whilst taking a swab sample from the greenhouse roof. The surrounding area harboured a low concentration of tomato production.

### Case study 6 (United Kingdom)

In March 2023, swab samples were collected from different surfaces of a ToBRFV experimental greenhouse in the North-East of England, UK (Fig 1c) where several ToBRFV inoculation experiments had taken place, including a ToBRFV-tomato sampling trial (Skelton et al, 2023b), followed by smaller ToBRFV experiments with infected plants. After the initial tomato trials, the greenhouse had been disinfected several times with Virkon (SLS Ltd, UK). The greenhouse had not contained ToBRFV-infected plants for at least one month before sampling. Swab samples were taken pre and post treatment with a hydrogen peroxide-based fogging product (BioDecon, UK). The surrounding area harboured a low concentration of tomato production, with the nearest producer situated over 40 km away.

### Sampling details

Case study 1: The cotton bud of a sterile swab was moistened by dipping it into a 1.5 mL tube with 150 μL DNAse/RNAse free water. The moistened bud was used to swab approximately 5 times across the target surface for a length of 5-10 cm while slightly rotating the stick after each swab. The bud was stored in the same 1.5 mL tube by snapping off the stick. A leaf sample consisted of leaf pieces of approximately 1-2 cm^2^ from 10 plants of the same species, or from an equivalent number of different leaves if less plants were present. Other samples included a piece of soil of approximately 1 cm^3^ collected from below the plastic covers on the greenhouse floor, a similarly sized piece of rockwool used for plant cultivation, or an approximately 2 cm^2^ piece of administration paper. Approximately 5 mL of wastewater, rainwater, or disinfected water was collected in tubes. Samples were collected in biological duplicates, by taking two swabs at the same time or dividing collected material in two equal parts or volumes.

Case study 2: Swab samples were taken by wetting the white cotton head of the swab (EUROTUBO® Collection swab, deltalab) in 1 mL sterile PBS (pH 7.4) and moving it back and forth over the surface of interest. Leaf samples were taken from 10 different tomato plants in 10 different rows spread randomly throughout the greenhouse for each location.

Greenhouse drain water was sampled from a collection reservoir awaiting disinfection, as described by Mehle et al. (2023).

Case study 3: Each sample consisted of leaf material from one plant per species.

Case study 4: Tomato leaf samples were taken as described for case study 2. At the time of sampling the tomato plants at location 1 were close to the end of the cropping cycle, whereas the plants at location 2 were younger, closer to the beginning of the cropping cycle.

Case study 5: Swab samples were taken by soaking cotton buds in 1 mL of 0.06 M phosphate buffer (pH 7) in a 50 mL universal tube. The soaked cotton bud was then used to swab approximately 20 cm^2^ of target surface, returned to the universal tube and soaked for at least 30 minutes. In addition, symptomatic leaf material, approximately 4 cm^2^, was taken from the growing crop.

Case study 6: Swab samples were taken from different greenhouse surfaces (door handle, glass pane, concrete floor, steel bench, aluminium ladder and watering lance), both pre and post treatment, with a hydrogen peroxide-based fog product at 7.5% active ingredient (approximately 40 min treatment).

In all case studies care was taken to avoid cross contamination and gloves were changed when taking different samples. Samples were stored at −20°C (case study 1) or 4°C (case studies 2-6) until RNA extraction.

### RNA extraction and (real-time) RT-PCR

Case study 1: 700 μL of guanidine hydrochloride (GH+) buffer (EPPO, 2022) was added to each swab sample and vortexed for 15 sec. Leaf, paper, soil, and rockwool samples were homogenized with GH+ buffer at an approximate ratio of 1:2-1:5 [w/v]. Each water sample was mixed with GH+ buffer at a ratio of 1:10 [v/v]. RNA was extracted from 500 µL of homogenate using the Qiagen RNeasy Plant Mini kit (manufacturer’s instructions). Real-time RT-PCR for ToBRFV was performed based on the protocol of the International Seed Health Initiative for Vegetable Seeds using the CaTa28 and CSP1352 primers and probes (ISF, 2020, EPPO, 2022) with two modifications: the TaqMan RNA-to-CT 1-step Kit (Applied Biosystems) was used instead of the Ultraplex 1-step Toughmix (Quantabio) and 2 μl of total RNA was used instead of 5 μl. The plant gene NADH dehydrogenase subunit 5 (nad5) was used as an internal control for RNA extraction from leaf samples (Botermans *et al*., 2013). Additional negative controls included cucumber and tomato leaves from young non-infected plants and clean printer paper.

Case study 2: RNA was extracted from 100 mg leaf or sepal tissue or from 300 µL of swab samples in PBS, using the RNeasy Plus kit (Qiagen). Swab samples were first thoroughly vortexed to stimulate release of the viral particles. Samples were analyzed by real-time RT-PCR for ToBRFV using the CaTa28 and CSP1352 primers and probes (ISF, 2020, EPPO, 2022)

Case study 3: RNA was extracted from leaf samples using the KingFisher Apex system (ThermoFisher) with the MagMAX^TM^ Plant RNA isolation kit (ThermoFisher). Samples were further analyzed by real-time RT-PCR for ToBRFV using the CaTa28 and CSP1352 primers and probes (ISF, 2020, EPPO, 2022). Additionally, RNA of the *S. oleraceus* and *G. quadriradiata* samples was used to perform a ToBRFV-specific near-full genome PCR (hereafter referred to as NFG-PCR) to determine the integrity of the detected RNA (Mehle et al., 2023), using primers F-22 and R-6392 as previously described by Eldan et al. (2022).

Case study 4: Samples were extracted and analyzed as described for leaf samples in case study 2 to confirm infection with ToBRFV.

Case Study 5 and 6: RNA was extracted from the swab buffer using the Qiagen RNeasy plant mini kit (manufacturer’s instructions). Nucleic acid was extracted from leaf material using GH+ buffer at 1:10 dilution. Leaves were ground using a HOMEX 6 (Bioreba) and RNA was extracted by magnetic bead extraction using Invimag Virus DNA/RNA mini-kit (Invitek GmbH). RNA extracts were tested by real-time RT-PCR with the Menzel and Winter primers (Menzel & Winter, 2021, EPPO, 2022) using iTaq universal probes one-step reaction mix (Bio-Rad) and 1 µl of RNA extract. Samples were tested in duplicate and Cq-values recorded. Cytochrome oxidase (COX) was used as an internal control for RNA extraction from leaf samples (Weller *et al*., 2000).

Appropriate positive and negative (extraction) controls were used in all case studies and performed as expected. Samples were considered positive when amplification curves were observed.

### Bioassay

Case study 1: Mechanical inoculations and cultivation of test plants were performed as described by Verhoeven and Roenhorst (2000). Plants were assessed for disease symptoms twice a week, and young non-inoculated leaves were tested by real-time RT-PCR 2-3 weeks after inoculation. Twelve biological duplicates of samples with relatively low Cq-values were used to inoculate three *Nicoticana benthamiana* and three *N. glutinosa* plants each.

Case study 3: Leaf material of *G. quadriradiata* and *S. oleraceus* was used to mechanically inoculate *N. tabacum cv Xanthi*. Inoculum was made by homogenizing 1 g leaf material in 3 mL 0.1 M phosphate buffer (pH 7.8) using metallic beads. Test plant leaves were dusted with carborundum (0.105 mm, VWR Chemicals) and 2 leaves of 3 plants per sample were inoculated, after which leaves were rinsed with tap water. The experiment was performed in a climate cell (PHCbi MLR-352H) under controlled conditions: 23°C (+/− 2°C), 10 h light period and 19°C (+/− 2°C), 14h dark period and repeated twice.

Case study 6: Swab samples were inoculated onto a *Nicotiana tabacum* plant as described by Skelton *et al*. (2023a).

Appropriate positive and negative inoculation controls were used in all case studies and performed as expected.

### Sequence analysis

Case study 4: The samples were processed using the Novel Enrichment Technique of Viromes protocol (NetoVIR protocol (Conceicao-Neto *et al*., 2015)), libraries were prepared using the NexteraXT Library Preparation Kit (Illumina). Sequencing was performed using the NextSeq500 platform (Illumina) for the leaf sample from location 1 and the NovaSeq6000 platform (Illumina) for the sample from location 2. The assigned sequencing clusters were adjusted to achieve similar sequencing depth in both platforms, approximately ten million reads per sample. The ViPER bioinformatics pipeline was used to process the sequencing data (De Coninck, 2021). The reads were quality trimmed using Trimmomatic (Bolger *et al*., 2014) and *de novo* assembly was performed using metaSPAdes (Langmead & Salzberg, 2012). The assembled contigs were classified using DIAMOND (Buchfink *et al*., 2015) and KronaTools (Ondov *et al*., 2011) with the lowest common ancestor approach. The two nucleotide sequences were aligned using MAFFT v7.453 (Katoh & Standley, 2013) and visualized using an in-house R script. Phylogenetic analyses were performed using the maximum likelihood method with TPM2u+F+G4 model using IQ-TREE (Nguyen *et al*., 2015) with 1000 bootstrap replicates.

## Results

### Case study 1: ToBRFV detection in and around nonhost production greenhouses in the Netherlands

Case study 1 aimed to gain insight in the presence of ToBRFV at locations with and without prior ToBRFV outbreaks and in areas with high and low concentrations of tomato production. Samples were obtained within and around ToBRFV nonhost (cucumber or green bean) production greenhouses from areas with a high concentration of tomato production (Western area) and a low concentration of tomato production (Eastern area). Two greenhouse types were studied: those that switched from tomato to ToBRFV nonhost production in the past six months but were still under official ToBRFV control measures (Western area), and those that had not grown tomatoes for at least the past 4.5 years (Western and Eastern area). In the Western area, three companies (no. 2-4) also cultivated tomatoes in separate compartments with additional hygiene measures taken by the company to prevent cross-contamination between compartments. For two of those companies (no. 2 and 3), a few samples were also collected from the tomato compartments for comparison.

### Western area

In the Western area with a high concentration of tomato production, ToBRFV was detected in multiple samples from all ToBRFV nonhost greenhouses as well as in one or more samples outside (Fig 2). Overall, ToBRFV positive samples from cucumber compartments of companies 1-4 with a ToBRFV outbreak in a recent previous tomato cultivation or with a current tomato cultivation in a nearby compartment showed lower Cq-values (Cq 13-39, mean Cq 24) than samples from companies 5-7 without recent tomato cultivation (Cq 23-31, mean Cq 29) (Fig 2a,c). However, Cq-values from company 1 (Cq 29-39, mean Cq 31) were relatively high, indicating a lower virus amount, and ToBRFV was detected in a lower number of samples than for companies 2-4 which were still producing tomatoes in different compartments. Cq-values from company 1 were more comparable to those of companies 5-7 which had not grown tomato in the past years. To assess whether there was a ToBRFV outbreak in the nearby tomato compartments of company 2 and 3, a few samples from young tomato leaves were tested and showed very low Cq-values (Cq 3), indicating an infection. Swab, soil, and water samples from the tomato compartments showed Cq-values comparable to those from cucumber compartments of the same companies (Fig 2b). In addition, a few samples were taken at three outside locations: around a hotel housing greenhouse staff, outside company 5, and across the road from a waste factory handling greenhouse waste. One or more positive samples were found at each location (Cq 21-39, mean Cq 29), including swabs from surfaces that are typically untouched (Fig 2d). Furthermore, ToBRFV was detected from multiple swabbed surfaces that were exposed to contact with individuals, plants, and objects in all sampled compartments, which might be attributed to contact transmission. However, ToBRFV positive samples were also detected from untouched surfaces at companies 2, 3, 4, 5 and 6. For instance, the ceiling frames several meters above the mature plants tested positive for companies 2, 3, 4 and 6 (Fig 2a-c). This indicated that ToBRFV residues also spread independently of contact, potentially through aerosols. The outside of the glass ceiling panes yielded positive ToBRFV test results for companies 2 and 3 (Fig 2a-b), whereas similar surfaces tested negative for the other companies, potentially due to rainfall before sampling which might have washed away virus residues. Interestingly, the inside surfaces of the glass ceiling panes consistently tested negative, possibly due to a lack of dust accumulation on the lower side of the ceiling panes.

**Figure 2.**
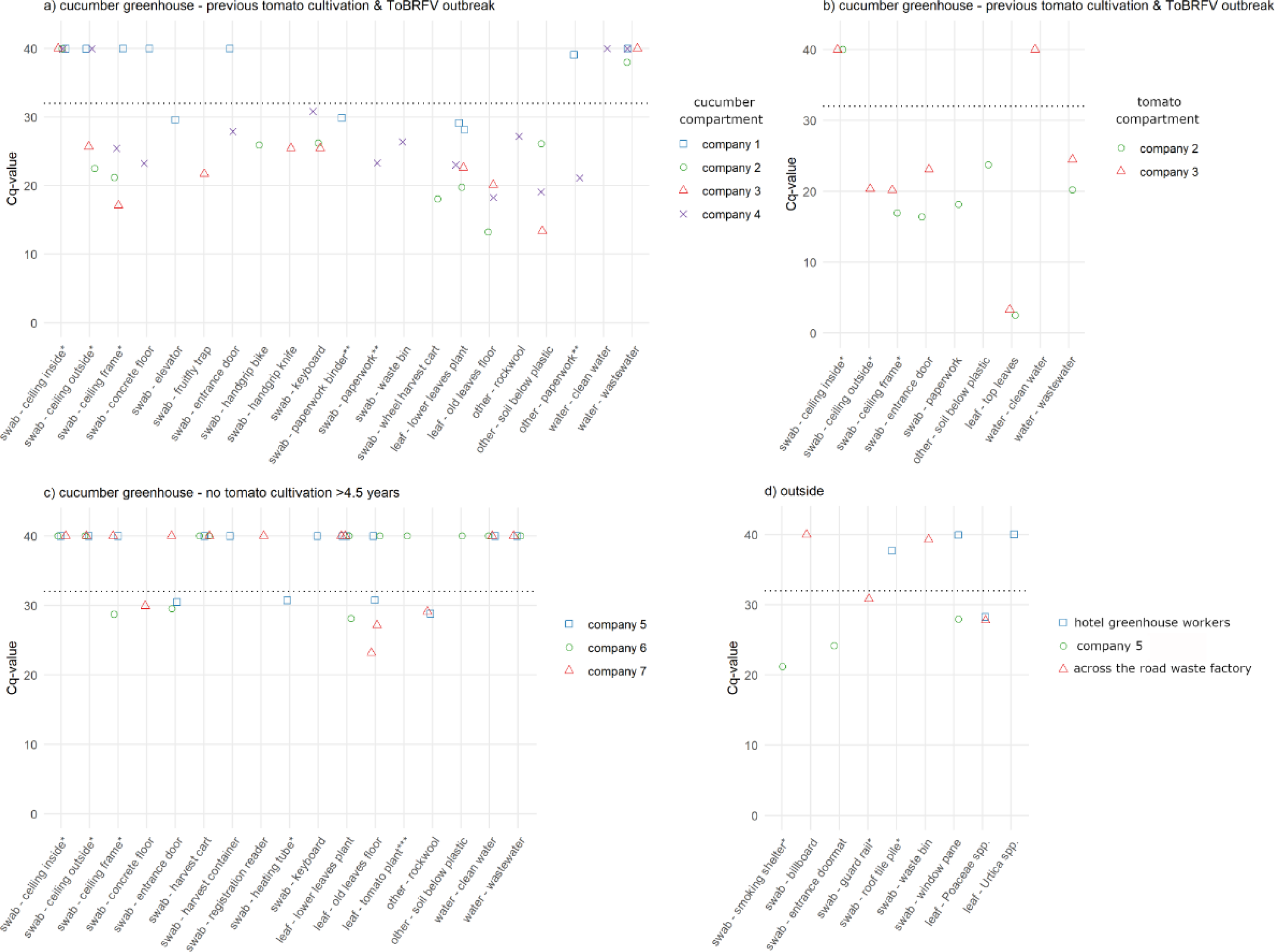
Cq-values of samples from the area with a high concentration of tomato production (Western Netherlands). a) Cucumber compartment of greenhouses with a previous ToBRFV outbreak, that switched to cucumber cultivation in all or some compartments. Companies 2-4 were still producing tomatoes in a different compartment. b) Tomato compartments of company 2 and 3, samples were taken for comparison with the cucumber (nonhost) compartments c) Greenhouses with no previous tomato cultivation in at least the past 4.5 years. d) Outside locations. Dotted line: threshold below which samples are considered positive for official samples in the Netherlands. *Surfaces typically untouched by indiviuals, plants and objects. ** Paperwork from the ToBRFV outbreak when tomato was grown, sampled in a room next to the cucumber compartment. ***A young volunteer tomato plant at the edge of the plastic covering the soil at company 6. Cq 40: no amplification. Cq-values refer to the CaTa28 primers and probes, comparable results were obtained for CSP1325 and controls reacted as expected. Points were plotted with a jitter function to prevent overlap.

At the cucumber producing companies with a previous ToBRFV outbreak, ToBRFV was detected on all leaf samples, both from leaves attached to living plants as from old leaves on the floor (Fig 2a). All soil and rockwool samples tested positive as well. In contrast, at companies without tomato cultivation in the past few years, only 4/12 cucumber leaf samples and 2/3 soil and rockwool samples were ToBRFV positive (Fig 2c). For the water samples, the only positive results were found for two wastewater samples from the tomato compartments that had not yet undergone water disinfection treatment (Fig 2b).

### Eastern area

In the Eastern area with a low concentration of tomato production, ToBRFV was detected with relatively high Cq-values of at least Cq 28 in only a few samples from cucumber and green bean production greenhouses (Fig 3a). At company 10, five hobby tomato plants were grown in between the cucumber plants, of which leaf samples tested negative. Outside company 10, leaves of three plant species, *Juglans regia, Pyrus communis* and *Poaceae* species tested positive but with relatively high Cq-values of 24-32. All other samples taken outside showed no amplification or Cq-values >32 (Fig. 3b). These results indicated that there was less environmental ToBRFV contamination in the Eastern area than in the Western area.

**Figure 3.**
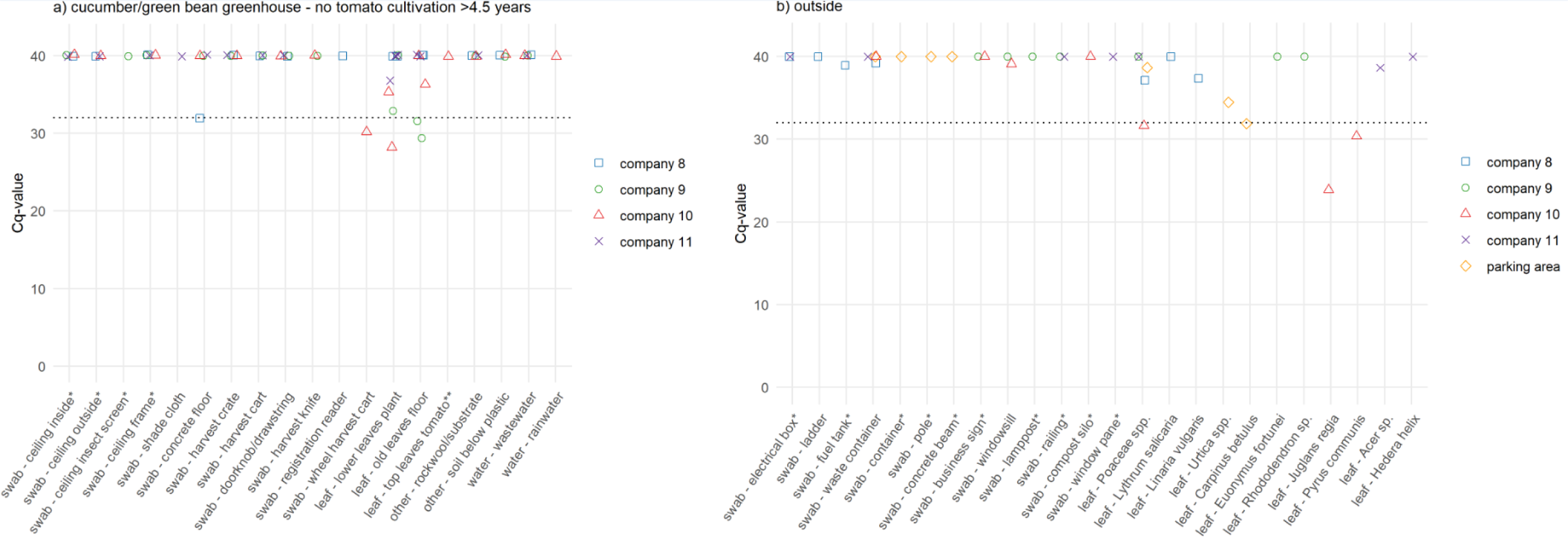
Cq-values of samples from the area with a low concentration of tomato production (Eastern Netherlands). a) Cucumber or green bean compartment of greenhouses with no previous tomato cultivation in at least the past 4.5 years. b) Outside locations. Dotted line: threshold below which samples are considered positive for official samples in the Netherlands. *Surfaces typically untouched by individuals, plants and objects. ** five hobby tomato plants grown in between the cucumber plants. Cq 40: no amplification. Cq-values refer to the CaTa28 primers and probes, comparable results were obtained for CSP1325 and controls reacted as expected. Points were plotted with a jitter function to prevent overlap.

To assess infectiousness, biological duplicates of 4 leaf, 5 swab, and 3 soil samples with relatively low Cq-values (tomato leaf: Cq 3, other samples: Cq 13-24) were inoculated onto *N. benthamiana* and *N. glutinosa*. Samples originated both from cucumber and tomato compartments in the Western Netherlands. Test plants only showed symptoms upon inoculation by the tomato leaf sample and presence of ToBRFV was confirmed by real-time RT-PCR (Table S1).

### Case study 2: ToBRFV detection in tomato production greenhouses at different outbreak stages in the Netherlands

To determine the infection status of three adjoining tomato production greenhouse units belonging to the same company, leaf and sepal samples were tested for ToBRFV. Results indicated that the three locations were in different phases of ToBRFV infection at the time of sampling (Table 2). At location 1, a ToBRFV outbreak was present as the leaf and drain water samples showed Cq-values of 6.5 and 22.9, and ToBRFV symptoms were observed in the crop. In contrast, at location 2, sepal and water samples showed Cq-values around 33 and no symptoms were observed. For location 3, sepal and water samples showed Cq-values of 23-24, suggesting the potential start of an outbreak at the time of sampling, although no symptoms were observed. To our knowledge, none of the sampled locations had been previously infected with ToBRFV.

**Table 2.** Detection of ToBRFV in tomato leaf and sepal and water samples of different greenhouse entities.

| Location | Sample type | Cq-value<sup>a</sup> | Symptoms observed |
| --- | --- | --- | --- |
| 1 | Leaf | 6.5 | Yes<sup>b</sup> |
|  | Water | 22.9 |  |
| 2 | Sepal | 33.4 | No |
|  | Water | 33.3 |  |
| 3 | Sepal | 22.9 | No |
|  | Water | 23.9 |  |

<sup>a</sup> Mean Cq-value obtained with CaTa28 and CSP1352 primers and probes.
<sup>b</sup> Symptoms involved mosaic and deformation in the head of the plant, and reduced leaf biomass; only observed in a couple of rows at the time of sampling.

Swab samples were collected from different surfaces, with varying degrees of contact with individuals, tomato plants, or objects, to determine the overall presence of ToBRFV at each location (Table 3). The lowest Cq-values from swab samples were obtained from location 1 (Cq 12 – 36, mean Cq 21.8), followed by location 3 (Cq 19 – 36, mean Cq 28.3). At these locations, ToBRFV was found on several surfaces that are typically untouched by individuals, tomato plants, or objects, such as the lights (mean Cq 23.3), fan (mean Cq 21.4) (location 1), and netting over the windows at the outside of the greenhouse (mean Cq 27.5) (location 3). While ToBRFV was detected in swab samples from the smoking rooms (Cq 21-36, mean Cq 29), cigarette butts showed higher Cq-values (Cq 35-36, mean Cq 35), suggesting that cigarettes were not the source of ToBRFV contamination in these rooms. The highest Cq-values were obtained from location 2 (Cq 24 – undetermined, mean Cq 34.6). Although Cq-values of tomato leaf samples from location 2 did not indicate an outbreak (Table 2), ToBRFV was still detected on several surfaces, most notably on the concrete path (mean Cq 31.3), groundsheet (mean Cq 30.9) and gutters (mean Cq 24.8). Overall, these results showed that ToBRFV was detected at all three locations, even when plants had no symptoms and produced high Cq-values.

**Table 3.**
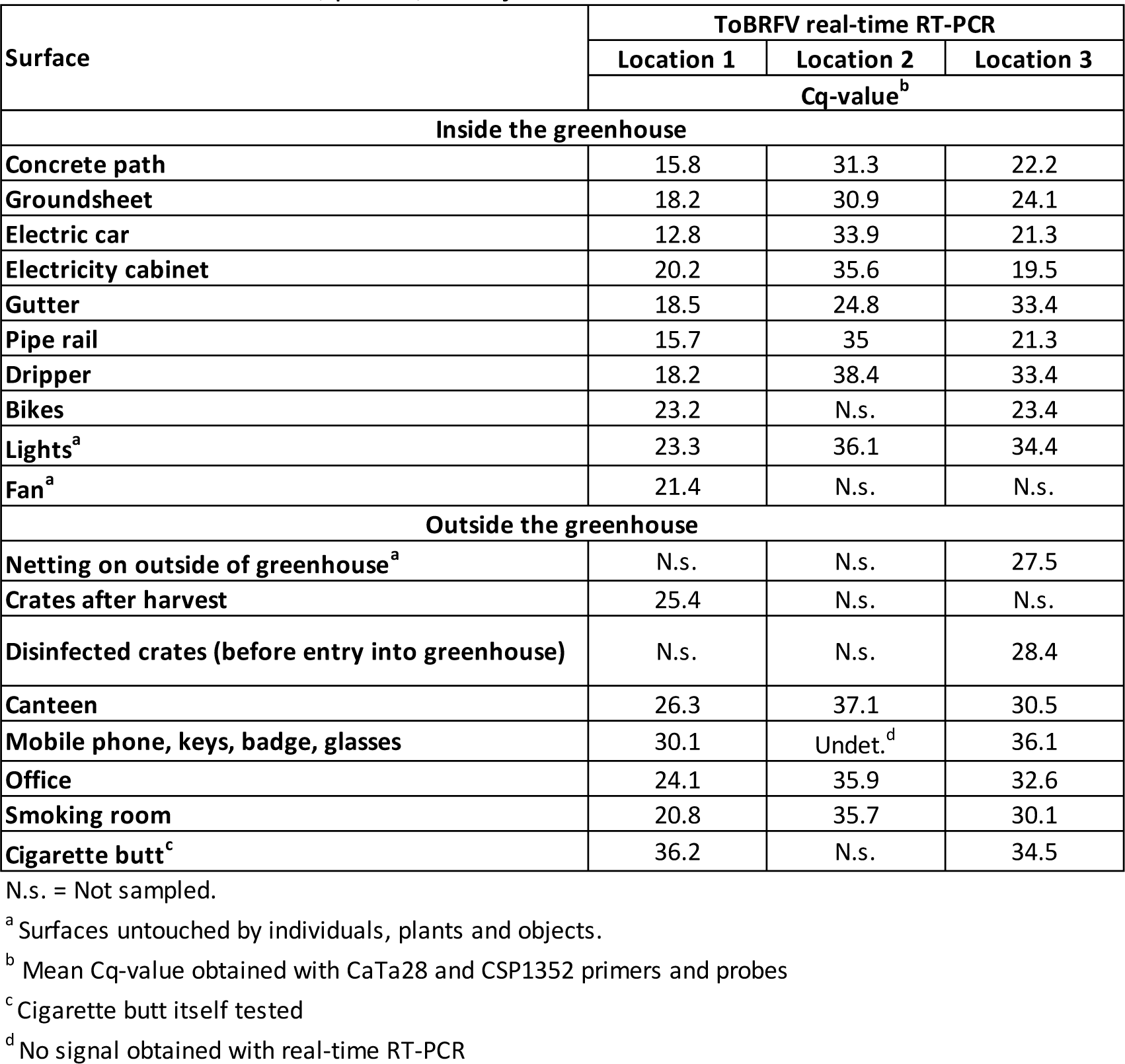
Results of swab samples taken from different surfaces with varying degrees of contact with individuals, plants, or objects.

### Case study 3: ToBRFV detection on weeds outside tomato production greenhouses in Belgium

To assess the presence of ToBRFV (residue) outside greenhouses, four weed species common to Western Europe were sampled in the North of Belgium in an area with a high concentration of tomato production. Samples were collected outside in the area immediately surrounding a tomato production greenhouse with a likely ToBRFV outbreak. ToBRFV was detected in all samples, with mean Cq-values ranging from 14.5 – 26.9 (Table 4). Cq-values (mean Cq 14.5 – 19.5) were lower for weeds that were sampled directly next to the greenhouse (touching the outside of the greenhouse), in comparison to weeds that were sampled at a distance of approximately 3 m (mean Cq 24.9 – 26.9). Mechanical inoculation leaf material from *G. quadriradiata* and *S. oleraceus* onto the local lesion host *N. tabacum L. cv Xanthi* did not lead to the appearance of lesions. Moreover, ToBRFV was not detected with the ToBRFV-specific NFG-PCR in these two weed samples, indicating that the positive real-time RT-PCR results were due to fragmented RNA.

**Table 4.** Cq-values of weed leaf samples taken outside a tomato production greenhouse in an area with a high concentration of tomato production in Belgium.

| Sample | Cq-value<sup>a</sup> | ToBRFV NFG-PCR |
| --- | --- | --- |
| *Sonchus oleraceus* | 14.5 | - |
| *Galinsoga quadriradiata* | 19.5 | - |
| *Solanum nigrum*<sup>b</sup> | 26.9 | n.t. |
| *Erigeron canadensis*<sup>b</sup> | 24.9 | n.t. |

:- No detection, n.t.: not tested
<sup>a</sup> Mean Cq-value obtained with CaTa28 and CSP1352 primers and probes
<sup>b</sup> Weeds sampled at approximately 3 m distance from the greenhouse (not touching the outside of the greenhouse)

### Case study 4: Transmission of ToBRFV between adjoining greenhouses in Belgium

In case study 4, the sequences of two ToBRFV isolates infecting tomato in two neighbouring greenhouse units of the same company were compared to investigate the possibility of transmission through viral residues. At month 0, the first isolate was obtained at location 1 from plants that started showing ToBRFV symptoms and on which ToBRFV was detected (Cq 11) (Table 5, Fig S1a). Three months after symptoms were first observed, the crop was cleaned out and the greenhouse was disinfected (Fig S1b). During crop clean-out, the infected plants were put outside to dry, close to location 2, where due to weather conditions some of the glass windows were open (Fig S1c). At month 3, the tomato plants at location 2 were not showing any symptoms and were therefore not tested for ToBRFV. At month 5, two months after th crop clean-out at location 1, the tomato crop at location 2 started showing symptoms and ToBRFV was detected (Cq 6) (Table 5, Fig S1d).

**Table 5.** Detection of ToBRFV in tomato leaf samples of two adjoining tomato production greenhouse units in Belgium.

| Location | Month | Cq-value<sup>a</sup> | Symptoms observed |
| --- | --- | --- | --- |
| 1 | 0 | 11.1 | Yes<sup>b</sup> |
| 2 | 5 | 5.8 | Yes<sup>b</sup> |

<sup>a</sup> Mean Cq-value obtained with CaTa28 and CSP1352 primers and probes.
<sup>b</sup> Symptoms observed involved yellow mosaic, deformation, and needle-tips on the leaves. No symptoms were observed on the fruits.

Subsequent HTS analysis produced two nearly complete ToBRFV genomes of 6371 nt and 6377 nt (GenBank accession no. OQ633211 and OQ633212). Both sequences contained four open reading frames encoding the viral small replicase subunit, RNA-dependent RNA polymerase (RdRp), Movement Protein (MP), and Coat Protein (CP). Phylogenetic analysis of the protein coding sequences indicated that these two isolates were 100% identical, but differed from publicly available ToBRFV sequences, (Fig S2). These results provide additional evidence that ToBRFV from location 1 served as the source of infection of the crop at location 2, presumably through ToBRFV residues.

### Case study 5: ToBRFV detection in a tomato production greenhouse in the UK

To investigate the detection of ToBRFV following a ToBRFV outbreak, swab samples were taken from a range of different surfaces in a UK tomato production facility formed of two conjoined greenhouses. ToBRFV was detected on all surfaces (Table 6), including some which had not been in direct contact with individuals, plants, or objects, such as the top of greenhouse lights, the inside of the fan units, and the entrance of the bee boxes. Of the 14 surfaces swabbed, 11 (79%) had mean Cq-values of less than 26, e.g. the roof structure (Cq 17.4), the ceiling lights in greenhouse 2 (Cq 25.3), the inside of the fans in both greenhouses (Cq 17.5 and 22.1), and the entrances of bee boxes (Cq 22.9 and 17.0). All swab samples taken from different areas of the greenhouse had some level of ToBRFV detection, indicating the widespread presence of ToBRFV residue.

**Table 6.**
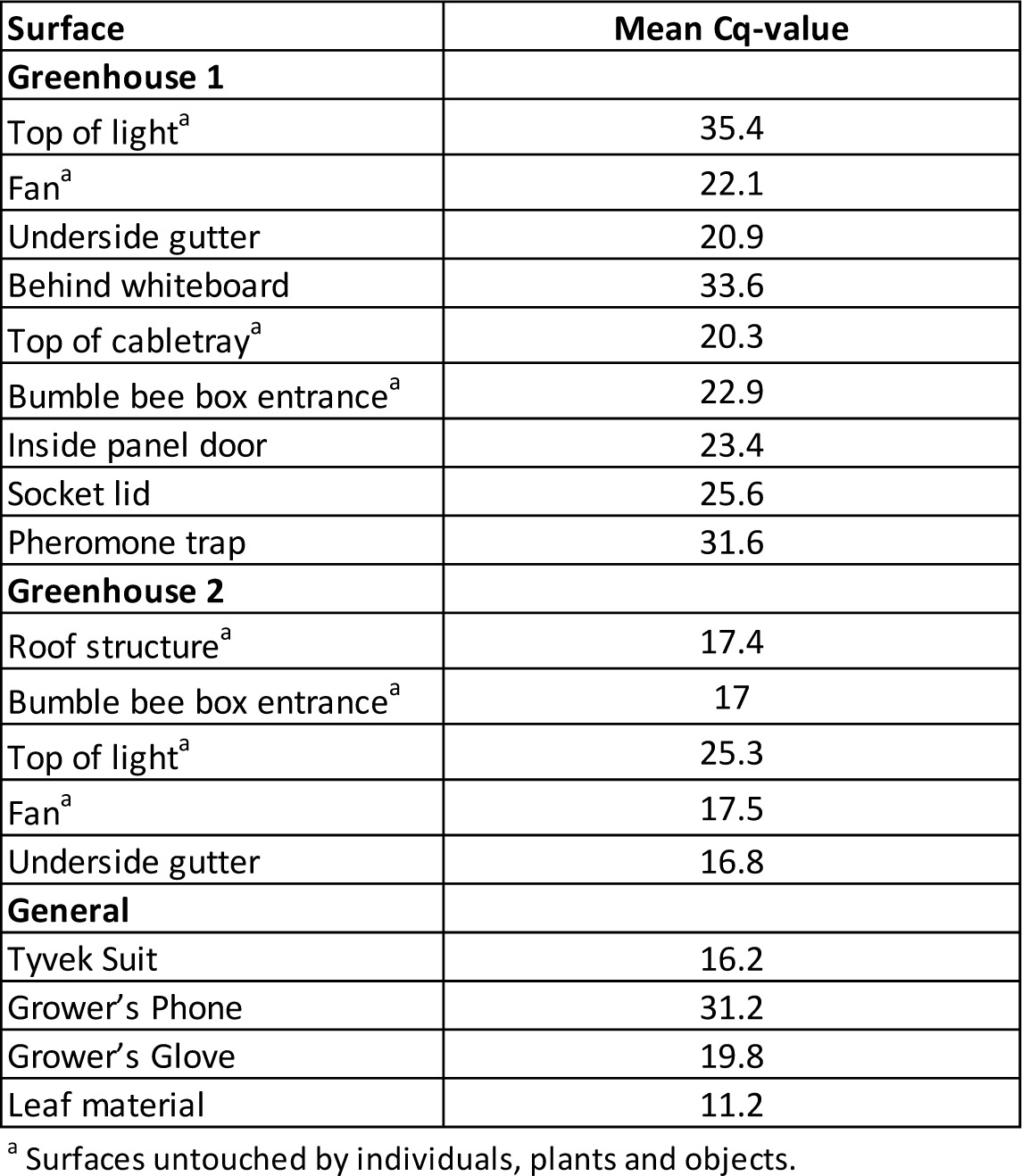
ToBRFV real-time RT-PCR results of swab samples taken from different surfaces with varying degrees of contact with individuals, plants, or objects at the UK tomato production greenhouse.

| Surface | Mean Cq-value |
| --- | --- |
| **Greenhouse 1** |  |
| Top of light<sup>a</sup> | 35.4 |
| Fan<sup>a</sup> | 22.1 |
| Underside gutter | 20.9 |
| Behind whiteboard | 33.6 |
| Top of cabletray<sup>a</sup> | 20.3 |
| Bumble bee box entrance<sup>a</sup> | 22.9 |
| Inside panel door | 23.4 |
| Socket lid | 25.6 |
| Pheromone trap | 31.6 |
| **Greenhouse 2** |  |
| Roof structure<sup>a</sup> | 17.4 |
| Bumble bee box entrance<sup>a</sup> | 17 |
| Top of light<sup>a</sup> | 25.3 |
| Fan<sup>a</sup> | 17.5 |
| Underside gutter | 16.8 |
| **General** |  |
| Tyvek Suit | 16.2 |
| Grower's Phone | 31.2 |
| Grower's Glove | 19.8 |
| Leaf material | 11.2 |

<sup>a</sup> Surfaces untouched by individuals, plants and objects.

The Tyvek suit, worn by the person taking swabs, sampled from where the person had leaned on the retaining bar on the scissor lift, had a mean Cq-value of 16.2. ToBRFV was also detected on the gloves worn by the grower (mean Cq-value 19.8). These were clean from the box when first put on, and no plant material was touched by the grower, but door handles and other equipment had been touched. The leaf material from the greenhouse had a mean Cq-value of 11.2 and could therefore have been a likely source of contamination.

### Case study 6: ToBRFV detection before and after disinfection in an experimental greenhouse in the UK

Swab samples were taken from different surfaces of a ToBRFV experimental quarantine greenhouse to assess the effect of disinfection treatment. ToBRFV was detected on all surfaces pre-hydrogen peroxide fogging treatment; mean Cq-values ranging from 19.8 (floor) to 28.5 (bench). Post fogging, the Cq-values from most of the surfaces increased, especially for the door and floor. Detection levels for the glass panes stayed mainly the same but the Cq-values for the bench decreased slightly post fogging treatment (Table 7).

**Table 7.**
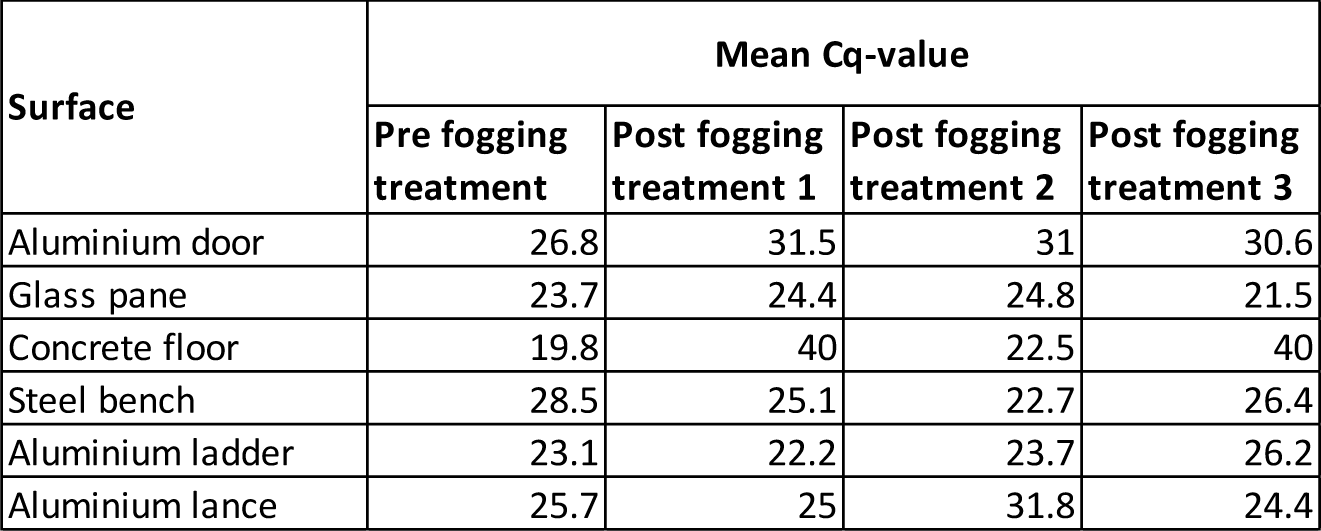
ToBRFV real-time RT-PCR results of swab samples taken from different surfaces of the UK experimental greenhouse, pre and post a hydrogen peroxide based fogging treatment.

Swabs were also taken from the surfaces before and after disinfection and tested by bioassay on *N. tabacum* to check for infectiousness. No virus symptoms were detected on any of the inoculated plants.

## Discussion

The case studies on ToBRFV presented here demonstrate the presence of detectable virus or residues in diverse types of samples and settings from sites in the Netherlands, Belgium, and the UK, both in the presence and absence of nearby infected plants. The detection of ToBRFV from nonhost plants and inanimate surfaces clearly shows the extent to which ToBRFV and/or its residues can spread beyond infected crops, highlighting the importance of contextual information for interpreting positive test results.

Previous studies have shown that tobamoviruses were detected in both the broader surroundings and in the immediate vicinity of crop production. Castello *et al*. (1995) detected infectious tobacco mosaic virus in fog and clouds in New York State and Maine, USA. In Slovenia, wastewater influent and effluent were shown to contain infectious pepper mild mottle virus, tobacco mild green mosaic virus, and tomato mosaic virus (Bacnik *et al*., 2020). Furthermore, in a commercial greenhouse with a high percentage of cucumber plants infected by cucumber green mottle mosaic virus, contamination of many surfaces has been shown after crop removal (Ellouze *et al*., 2020). Additionally, a study at a ToBRFV-infected tomato greenhouse in Germany showed that infectious ToBRFV was detected from swab samples of different surfaces not only in the greenhouse, but also in the packaging hall and staff accommodation, presumably as a result of cross contamination by human contact (Ehlers *et al*., 2023).

In addition to contact transmission, case studies 1, 2, and 5 indicated that ToBRFV residues can also spread independently of contact, as swabs from surfaces that are typically not in contact individuals, plants, or objects tested positive. Skelton et al. (2023b) hypothesized that erratic detection of ToBRFV from non-inoculated control plants was due to environmental contamination of the virus originating from highly infected plants in close proximity. This environmental residue contamination may have included aerosols carrying the virus or finely powdered dried material (dust) from infected plants. This hypothesis was supported by the observation that at ToBRFV outbreak sites, lower Cq-values were observed in samples from older leaves compared to younger leaves or sepals of asymptomatic plants, which might be explained by the accumulation of dust (Skelton et al., 2023b).

Furthermore, the detection of ToBRFV in greenhouses producing a nonhost crop, which had no prior history of tomato cultivation (case study 1), indicated that positive test results could not be attributed to residual virus from a previous cultivation, but rather must have originated from external sources. The often dusty outside of the glass ceiling panes tested positive in some cases, whereas the non-dusty inside of the glass ceiling panes never tested positive, not even from tomato compartments with an ongoing ToBRFV outbreak. This supports the hypothesis that ToBRFV (residue) was spread through aerosols, containing the virus or dried material from infected plants, such as leaf particles, pollen, and trichomes (dust). The airborne dispersion of virus or viral residues could also explain the detection of ToBRFV outside of greenhouses, as was found in case studies 1 and 3.

Proximity to a source of infection has been observed to positively affect the rate of ToBRFV spread before (Ehlers et al., 2023). Indeed, the highest levels of ToBRFV were found in samples from companies with a previous ToBRFV outbreak and adjacent tomato cultivation in case study 1. Similarly, in case study 2 the highest levels of ToBRFV surface contamination were found at the locations with the lowest Cq-values in tomato. In case study 3, weeds sampled closest to a greenhouse with a likely ToBRFV outbreak also showed the lowest Cq-values. For the companies without a previous ToBRFV outbreak or no adjacent tomato cultivation, ToBRFV was detected from fewer samples and with higher Cq-values (case study 1). Presumably, these positive samples were due to the presence of ToBRFV residues in the environment.

In case study 1, a high ToBRFV virus pressure was present at the time of sampling, since in 2023 ToBRFV was identified from many locations, predominantly in the Western part of the Netherlands (https://nextstrain.nrcnvwa.nl/ToBRFV/20240205). In addition, genomic analyses have indicated that at tomato production sites ToBRFV was presumably applied on purpose to plants, even though it is a quarantine pathogen (Botermans *et al*., 2023). The combination of purposeful application of a quarantine virus and subsequent natural spread may have resulted in a relatively high virus pressure in the Netherlands, especially in the Western area.

Indeed, case study 1 showed the detection of ToBRFV across all sampled locations in the Western Netherlands, both inside and outside greenhouses, indicating a widespread presence of the virus or virus residues in the region. In the Eastern Netherlands, with a much lower concentration of tomato production, ToBRFV was detected from fewer samples and with clearly higher Cq-values than in the Western area, suggesting that a lower concentration of inoculum in an area may lead to lower levels of environmental residues. Overall, the limited number of samples in this study hindered statistical comparisons between sample types and locations. However, there was a clear trend of more positive samples, with a higher viral load for those locations that were in close proximity to tomato production. Beyond the diagnostic risk of detecting ToBRFV in the absence of infected plants, a high virus presence can also oppose the effectiveness of novel ToBRFV-resistant cultivars, since this increases the likelihood of the emergence of ToBRFV isolates that can break through resistances. The first mutant that could break through a newly introduced ToBRFV-resistant cultivar has already been detected (Zisi *et al*.). Furthermore, the fact that ToBRFV resistant cultivars currently on the market do not provide complete immunity further highlights the importance of achieving and maintaining a low virus pressure.

Due to the persistent nature of tobamoviruses, however, it is difficult to eradicate these viruses as they can remain infectious under extreme conditions and for extended periods of time (EPPO, 2024). ToBRFV has been found to remain infectious for at least 28 days on some surfaces such as aluminum and stainless steel and up to six months for plastics and glass (Skelton et al 2023a). Such variations in the survival rate on different surfaces have also been observed for potato spindle tuber viroid (Mackie et al 2015). These extended survival times potentially enable environmental residues to act as an inoculum source of future crops.

To gain further insight in the potential transmission of ToBRFV from environmental residues, bioassays were performed with biological duplicates of several nonhost leaf, swab, and soil samples from case study 1, exhibiting relatively low Cq-values. None of the inoculations, however, resulted in virus transmission, neither did the inoculations from weed samples in case study 3, nor the swab samples from different surfaces in case study 6. In contrast, case study 4 suggested possible transmission of ToBRFV between two neighbouring greenhouses from the same company in Belgium. Based on the timing of infection in the two greenhouses and the high degree of sequence similarity among the respective virus isolates, it is likely that transmission happened through aerosols from dried infected plant material. However, it cannot be fully ruled out that transmission occurred via other routes (EPPO, 2024), but workers and tools were separated for both greenhouses and rigorous hygiene measures were applied, so it is considered unlikely that contact transmission played a role in transmission of the virus. This case study shows that infectiousness of viral residues cannot be excluded. Even a very low rate of virus transmission from the environment to plants may lead to widespread infection due to the ease of transmission from one infected plant to another. This emphasizes the necessity of collecting contextual information regarding infection status, infection history, and sampling location to assist in result interpretation.

The potential of transmission through environmental residues highlights the importance of effective ways to inactivate ToBRFV and other contact transmissible pathogens from surfaces following an outbreak, and the need for effective overarching disinfection protocols being used as part of eradication measures (Skelton et al., 2023a). Case study 6 showed that even after disinfection with hydrogen peroxide based fogging treatment, ToBRFV was detected from surfaces such as doors and windows, even though the virus appeared no longer infectious. These findings further support the previous reports of detection of ToBRFV from a range of substrates and surfaces following disinfection (Skelton et al., 2023a, Davino et al., 2020).

The detection of ToBRFV on diverse plant samples collected outside greenhouses, raises questions on their role in the epidemiology of this virus. In both studies in the Netherlands (case study 1) and Belgium (case study 3), plants gave positive test results in real-time RT-PCR and negative results in bioassay. In addition, the results of the ToBRFV-specific NFG-PCR tests in case study 3 indicated the absence of intact viral RNA. Given that the sampling in case study 3 occurred near a company likely experiencing an ongoing ToBRFV outbreak, it is plausible that the detection of ToBRFV on weeds could be attributed to the spread of virus residues from this company and does not prove infection. Recent literature suggested that the detection of ToBRFV from weeds indicates their natural host status (Cultrona *et al*., 2024, Salem *et al*., 2022a), although also in these studies it cannot be excluded that the detection of ToBRFV on weeds resulted from contamination with ToBRFV residues present in the environment. The host status of these species needs confirmation, and at present there are no indications that weeds play a role as a ToBRFV reservoir.

Regarding diagnostics, the increased use of sensitive methods like real-time RT-PCR, enables the detection of pathogens at relatively low concentrations. In the case of ToBRFV, however, positive test results may either indicate actual (nascent) infections or the presence of residual genetic material in the environment. The case studies here show the detection of ToBRFV in leaf samples from nonhost plants and other sample types, probably due to environmental ToBRFV residues, which raises the possibility that healthy, uninfected tomato plants also yield positive test results. This presents diagnostic challenges when dealing with a stable virus like ToBRFV, especially in areas with lots of environmental residues. Therefore, depending on the purpose of testing, Cq-values may need to be interpreted in conjunction with contextual information, such as the origin of the sample, symptoms observed, the probability of residues from prior cultivations and the likelihood of residues in the broader environment. This study highlights the complexity of decision-making in response to positive test results and underscores the need for a nuanced approach to manage the impact of ToBRFV in agricultural settings, together with further research to relate ToBRFV detection data to the risks of creating new infections.

## Acknowledgments

We thank our NVWA colleagues René van den Berg, Wietse den Hartog, Aron van Duijnhoven, Carla Oplaat, Gilo Pleunis, Karen van der Ven, Naomi te Braak, Stephanie Rensen, Susan Fekken, Tim Warbroek, Bart van de Vossenberg, Floyd Pere and Monique Slegers for their assistance with case study 1. We also thank the B2B (Beheersing van ToBRFV op de Belgische tomatenbedrijven) consortium (grant number HBC.2021.1075), including the Belgian research stations of Sint-Katelijne-Waver (PSKW) and Hoogstraten (PCH) for their help with case study 3. Finally, we thank the Flanders Innovation & Entrepreneurship (VLAIO) for the funding provided in the framework of Baekeland Mandate HBC.2020.2306 for case study 4. The UK case studies were funded under the Defra-Fera Long Term Service Agreement.

## Data availability statement

The nucleotide sequences from case study 4 can be found in GenBank (https://www.ncbi.nlm.nih.gov/genbank/) with GenBank accession no. OQ633211 and OQ633212.

## Supporting information

**Table S1.**
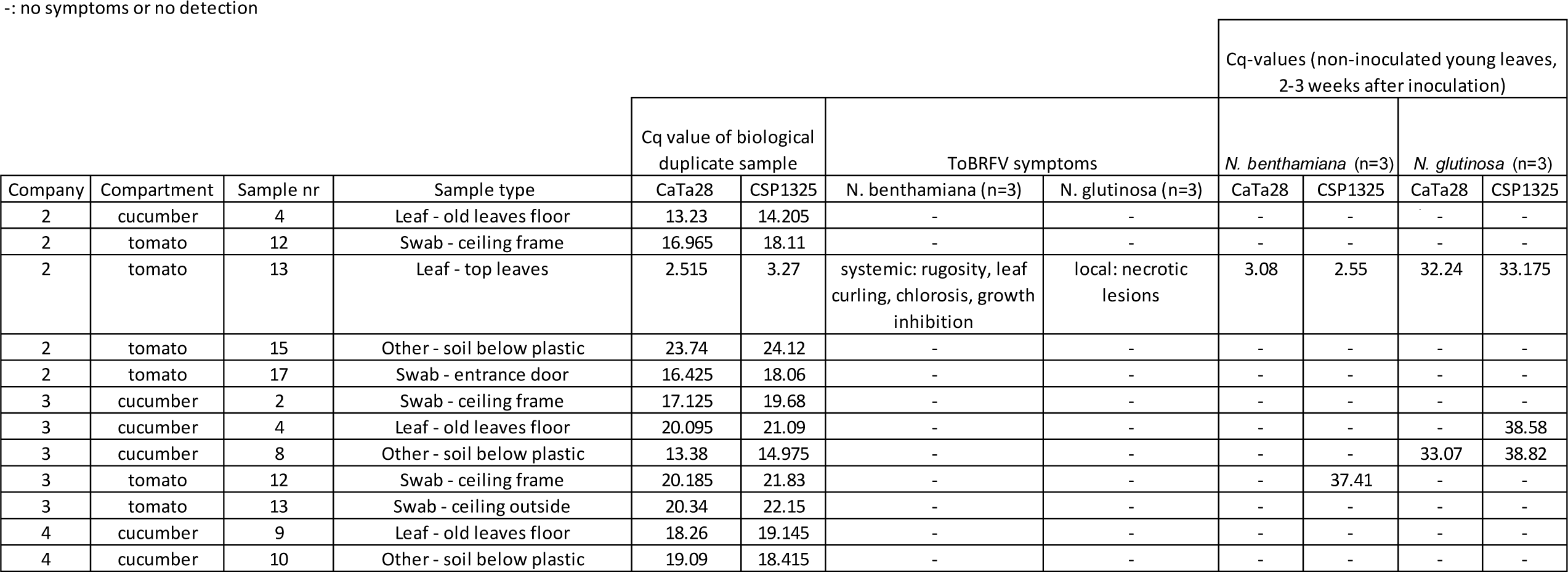
Symptoms and Cq-values on *Nicotiana benthamiana* and *N. glutinosa*, after inoculation with samples of which biological duplicates showed relatively low Cq-values (case study 1).

**Figure S1:**
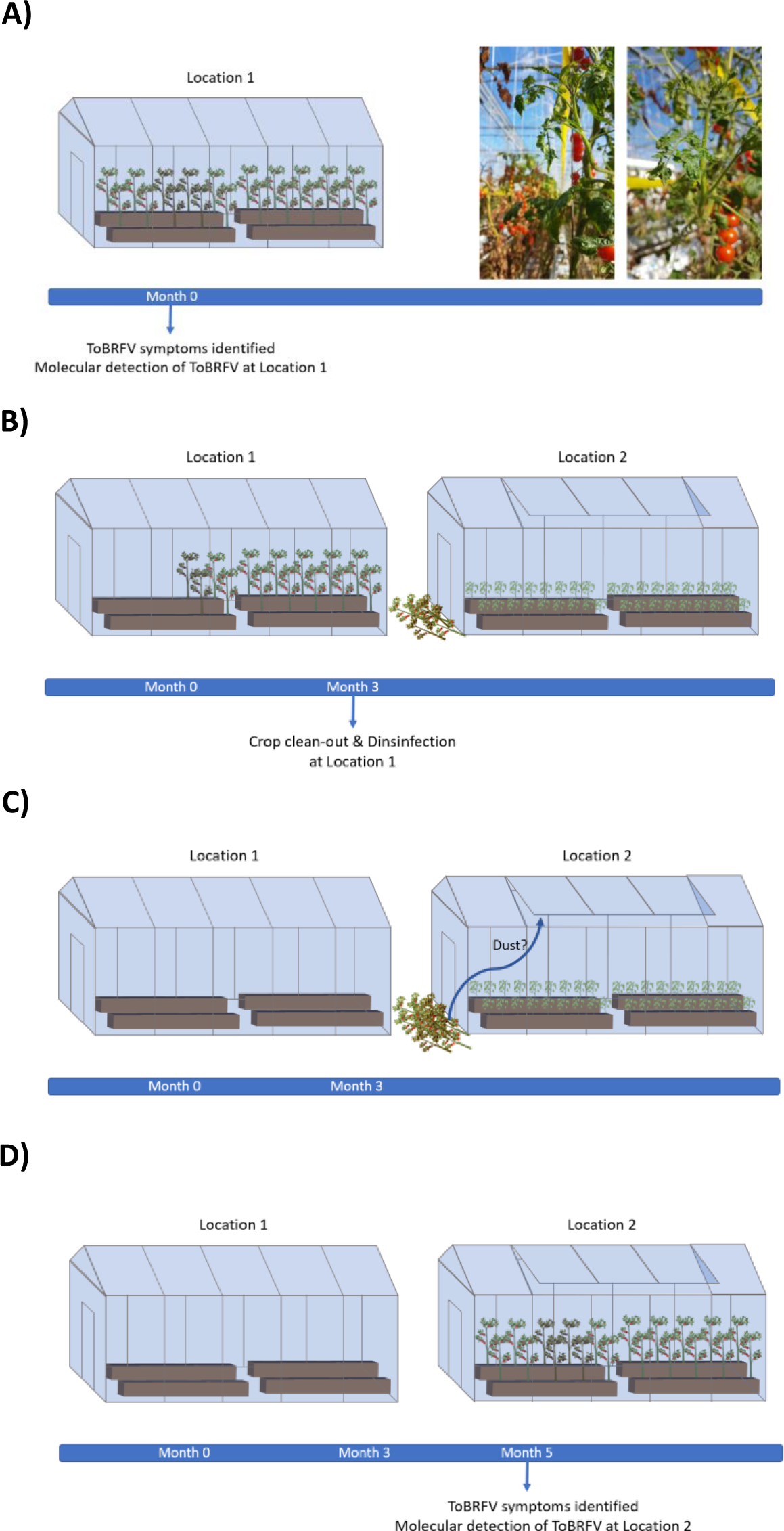
Hypothesis for transmission of ToBRFV through dust (case study 4). a) At location 1, ToBRFV symptoms were identified (Month 0). Infection was confirmed by real-time RT-PCR. b-c) Three months later, the ToBRFV-infected crop at location 1 was cleaned out and the greenhouse was disinfected. The cleaned-out crop was put outside to dry, close to location 2. Location 2 had opened some of its windows due to weather conditions, allowing dust from the decomposing infected crop of location 1 to enter the greenhouse. d) At month 5, ToBRFV symptoms were identified at location 2. Infection was confirmed by real-time RT-PCR. Figures were created with Biorender.com.

**Figure S2.**
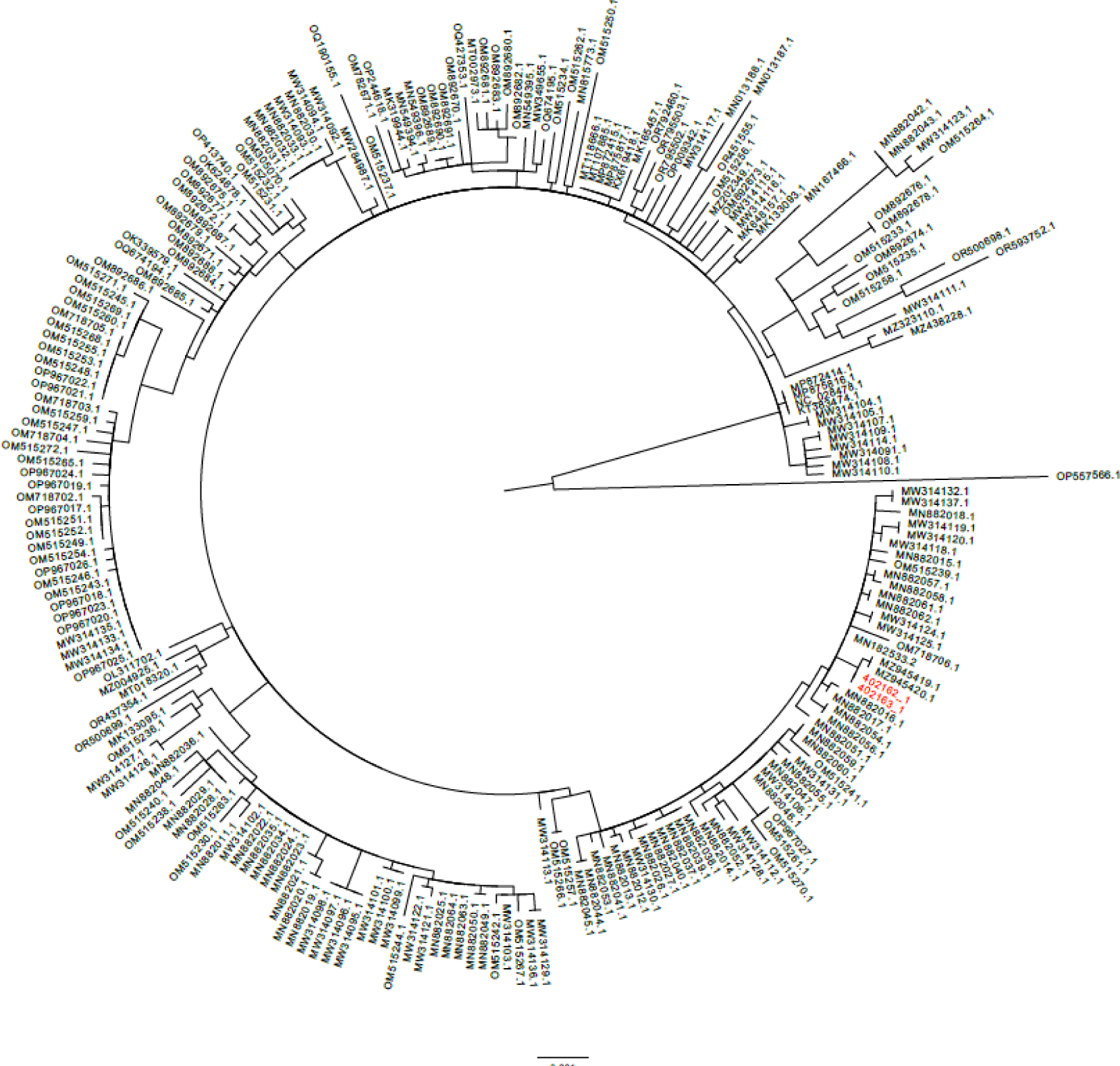
ToBRFV phylogenetic analysis based on the protein coding sequences (case study 4). Maximum-likelihood tree of the ToBRFV isolate sequences generated in this study are indicated in red (Isolate 402162_1 (GenBank accession no. OQ633211) and 402163_1, (GenBank accession no. OQ633212), whereas publicly available complete genome sequences from NCBI (accessed on 04/01/2024) are indicated in black.

